# Effects of aging on upper body express visuomotor responses while reaching under varying postural demands

**DOI:** 10.1101/2025.06.11.659162

**Authors:** Lucas S. Billen, Madeline Gilchrist, Vivian Weerdesteyn, Brian D. Corneil

**Affiliations:** Department of Rehabilitation - Donders Institute for Brain, Cognition & Behavior, Radboud University Medical Center, Nijmegen, NL; Department of Psychology, Western University, London, CA; Sint Maartenskliniek Research, Nijmegen, NL; Department of Physiology & Pharmacology, Western University, London, CA; Robarts Research Institute, London, CA

## Abstract

Humans can react remarkably quickly to novel or displaced visual stimuli when time is of the essence. Such movements are thought to be initiated by a subcortical fast visuomotor network, but it is unclear how this network declines with age. Past work in the upper limb has detailed delayed reaching corrections to jumped visual stimuli in the elderly, but the underlying mechanisms contributing to these changes of the fast visuomotor network are poorly understood. Conversely, work in the lower limb has reported delayed muscle recruitment during obstacle avoidance, but such findings may be confounded by age-related challenges in postural control. The output of the fast visuomotor network can be quantified by measuring express visuomotor responses (EVRs), which are the earliest and very short-latency bursts of muscle activity that follow visual target presentation. Here, we compare the prevalence, latency, and magnitude of EVRs in elderly (58-80 years old) and younger (18-25 years old) participants performing visually-guided reaches. We also investigated the impact of postural stability by having participants reach either while seated on a stable chair, or on a wobble stool. Both the elderly and younger cohorts expressed EVRs, but EVRs in the elderly were comparatively less frequent, and had longer latencies and smaller magnitude. Postural instability had no effects on these outcomes. Our results suggest age-related declines in the fast visuomotor network, potentially resulting from deterioration of underlying circuits and a prioritization of stability over speed. This study serves as an important standard for future research investigating clinical populations.

**Highlights:**

- Do aging and postural stability affect express visuomotor responses (EVRs)?
- EVRs were recorded from the pectoralis muscle during rapid goal-directed reaches
- EVRs were smaller and delayed, and movement was slower, in elderly participants
- Age-related differences were independent of our postural stability manipulation
- Our results suggest age-related declines of the fast visuomotor network

## Introduction

The ability to interact with our dynamic environment is essential. Rapid goal-directed movements towards visual stimuli are important both in reaching, for example when catching a ball thrown towards us, and in stepping, like when intercepting a ball while playing soccer. Declines in this ability with age may lead to an increased falling risk. Over the years, evidence has accumulated in support of the idea that a subcortical fast visuomotor network may initiate such rapid visuomotor transformations. For example, studies where a visual target is displaced during an ongoing movement have reported short-latency on-line reach corrections that are thought to be initiated subcortically (Day & Brown, 2001; Day & Lyon, 2000; Fautrelle, Ballay, et al., 2010; Fautrelle, Prablanc, et al., 2010; Mekhaiel et al., 2025). One way of assessing the output of this network, even for movements initiated from rest, is via electromyographic measurement of Express Visuomotor Responses (EVRs) (Corneil et al., 2004; Gu et al., 2016; Kozak et al., 2019; Pruszynski et al., 2010), which are the first detectable changes in muscle recruitment following presentation of a new or shifted visual target. These short-latency (within less than 100 ms of target onset) bursts of muscle activity facilitate movement toward the visual target, and are thought to originate in mid-brain superior colliculus, from where they are relayed to the motor periphery via the subcortical tecto-reticulospinal tract (Corneil & Munoz, 2014; Glover & Baker, 2019; Pruszynski et al., 2010).

Prominent in many studies on EVR expression is the hypothesis that, while EVRs are reflexively triggered by the visual stimulus, higher cortical areas can contextually modulate the amplitude of the network’s output. Indeed, various cognitive factors have been shown to influence EVR magnitudes, such as the predictability of the timing of the visual stimulus (Contemori et al., 2021; Wood et al., 2015) or the intention to reach towards or away from the presented target (Gu et al., 2016). We recently demonstrated that EVRs in the lower limb were readily expressed when stepping from a stable posture, but suppressed when stepping from an unstable posture (Billen et al 2023), indicating that the perception of postural (in)stability may similarly modulate EVR expression.

Interestingly, in an ensuing study in older individuals with and without Parkinson’s disease, the presence and magnitude of EVRs in the lower limb appeared to be lower and reaction times longer (Billen et al., 2024) compared to those previously observed in younger participants (Billen et al., 2023; Giesbers et al., 2025). While a direct comparison could not be made between young and elderly subjects due to methodological differences between the two studies, these observations suggest potential age-related degradation of the subcortical visuomotor network. Supporting this, muscle recruitment onsets are delayed in elderly participants during obstacle avoidance tasks (Weerdesteyn et al., 2007; Zhang et al., 2021), which is thought to involve the same visuomotor network. However, the stepping tasks used in these and the abovementioned studies (Billen et al., 2023, 2024) inherently involve postural control, which also declines with age (Bugnariu & Sveistrup, 2006; Maki & McIlroy, 1996). It therefore cannot be excluded that greater postural control difficulties in aging might have caused the attenuation of the fast visuomotor network. Indeed, previous findings in both healthy participants (Reynolds & Day, 2005) and, to a larger extent, in mildly affected stroke patients (Nonnekes et al., 2010) demonstrated that rapid stepping adjustments were aborted if these adjustments impacted balance, suggesting that the fast visuomotor network’s output may be suppressed in order to safely complete the step.

In the upper limb, multiple studies have demonstrated longer movement reaction times during online reach corrections in elderly participants (Kimura et al., 2015; O’Rielly & Ma-Wyatt, 2020). However, while these movement-related parameters quantify age-related changes in behavior, they do not provide a direct readout of the fast visuomotor network. Furthermore, it is unclear whether the fast visuomotor network is even engaged if latencies of corrective response exceed 200 ms, as these RTs are in the realm of voluntary reactions (Sarlegna, 2006). In contrast, EVRs, captured through surface EMG, provide a more objective measure of these rapid visuomotor transformations, but, to date, no study has investigated the effects of aging on the fast visuomotor network’s output in isolation. Therefore, we aim to directly compare EVR expression in younger and elderly participants in the upper limb during a seated reaching task - a movement that typically does not challenge postural control. Increasing postural instability in separate blocks of trials allows us to distinguish between the effects of aging and postural control on EVR expression. We hypothesized that reaching from a stable sitting posture would evoke robust EVRs in both groups, but with larger magnitudes in the younger participants compared to the elderly ones, which is in line with the indirect comparison made based on lower body EVRs (Billen et al., 2023, 2024). When increasing the instability of the sitting posture, we expected a decrease in EVR expression (both in prevalence and magnitudes) in both groups, with a more pronounced decrease in the elderly group. This would reflect the greater effort required to maintain balance, potentially reflecting prioritization of postural stability over rapid motor responses.

## Methods

### Subjects

22 young healthy subjects (16 females, 7 males; age range: 18-25 years (*M =* 18.9, *SD =* 2.2)) and 16 elderly healthy participants (8 females, 8 males; age range: 58-80 years (*M =* 68.8, *SD =* 6.7)) participated in this study. The younger participants were recruited through the Psychology Research Participant Pool. The elderly participants were recruited through the MacDonald Lab volunteer database at Western University. None of the participants had any visual, neurological, or motor-related disorders that could influence their performance in the study. The study protocol was approved by the Health Sciences Research Ethics Board of the University of Western Ontario (London, Ontario, Canada) and the study was conducted in accordance with the latest version of the Declaration of Helsinki. All participants provided written informed consent prior to participation and were free to withdraw from the study at any time. Participants received 40CAD as a monetary reward. Three participants from the elderly group were excluded: two due to low-quality EMG recordings and one because the experimental task proved too challenging.

### Apparatus & experimental task

Participants performed the experiment using a KINARM End-Point Lab (BKIN Technologies, Kingston, ON, Canada) and made reaching movements with their right arm (Figure 1A). Kinematic data were sampled at 1 kHz by the KINARM platform. Visual stimuli were projected onto an upward-facing mirror from a custom built-in projector (PROPixx projector by VPixx, Saint-Bruno, QC, Canada; custom integrated into the KINARM End-Point Lab). Direct vision of the hand was obscured by a shield beneath the mirror, but hand position was represented on the monitor in real time via a real-time cursor projected onto the screen. Surface EMG electrodes (Bagnoli-8 system, Delsys Inc., Boston, MA, USA) were used to record activity from the clavicular and sternal heads of the right pectoralis major, sampled at 1000Hz (Figure 1C). In line with previous studies (Kozak et al., 2020), a constant loading force of 5 N to the right and 2 N toward the participant was provided to increase the baseline activity of the muscle of interest, so that the EVR could be expressed as either an increase or a decrease in muscle activity following leftward or rightward target presentation, respectively. This constant load is fairly small and was well tolerated even by the elderly subjects.

**Figure 1.**
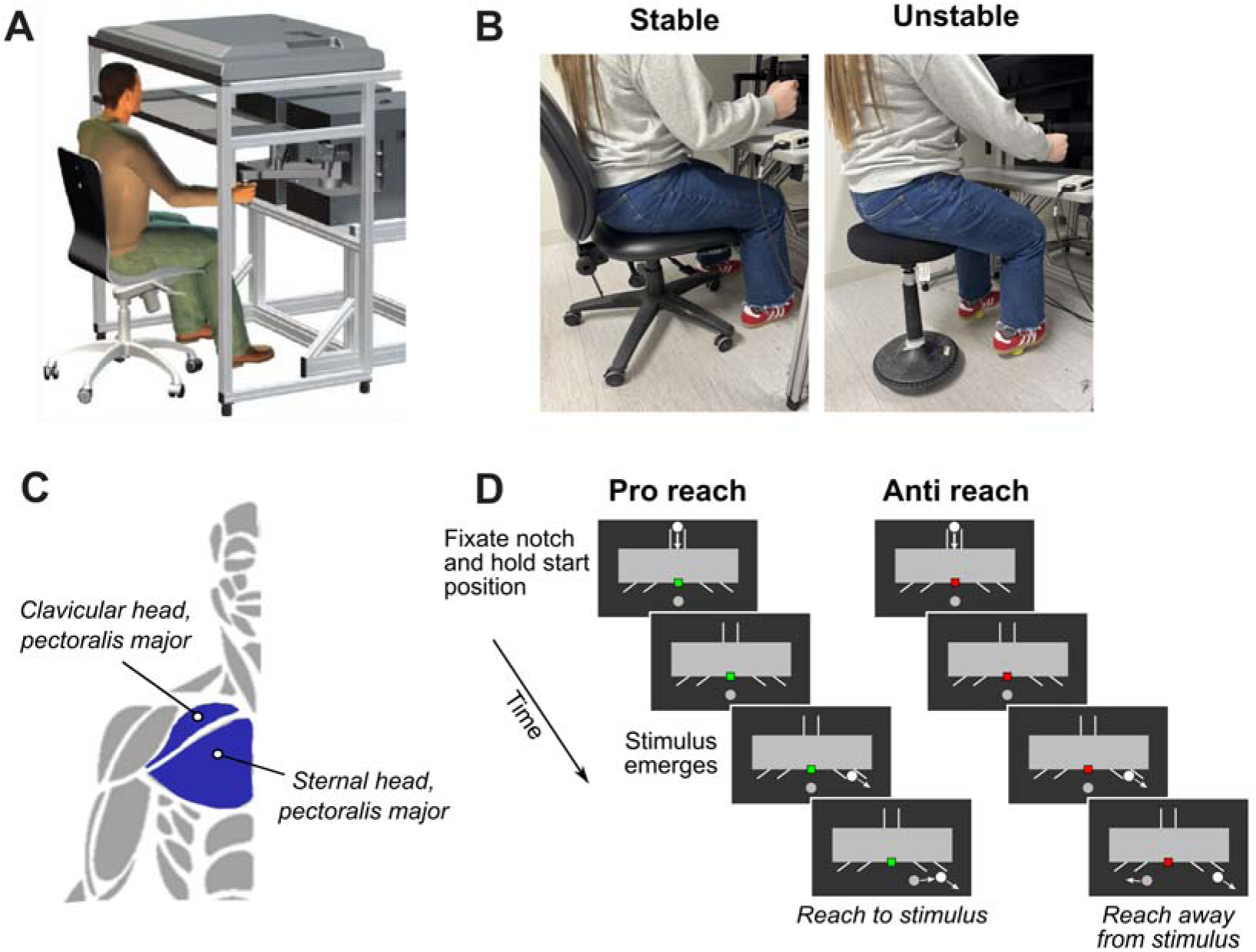
Behavioral paradigm. **A**: Participant moved the manipulandum of a Kinarm End-point robot with their right hand. **B**: Participants were either seated on a stable chair or on a wobble stool with a rounded base and with their feet placed on two tennis balls. **C:** Surface electromyographic recordings were taken from the clavicular and sternal heads of right pectorals major, which in this posture contributes to leftward movements of the right arm. **D**: We used a variant of the emerging target paradigm (Kozak et al., 2020). To start the trial, participants acquired the central start position (grey circle) and fixated a small notch at the bottom of the occluder. The colour of the notch indicated whether a pro-reach (green) or anti-reach (red) movement was to be performed. This instruction was available for the entire trial, and for at least 1500 ms before target emergence.

We used a variant of the emerging target paradigm that was previously described in Kozak et al. (2020) (see Figure 1D). Each trial started with the appearance of a stationary visual stimulus (solid white circle within a vertical channel), presented on a black background, that appeared above an occluder. At the center point of base of the occluder, a small rectangular instruction cue indicated whether participants were to respond to stimulus emergence with a reaching movement towards the stimulus (green rectangle, a ‘pro-reach’ trial) or away from the stimulus (red rectangle, an ‘anti-reach’ trial). This instruction cue was present throughout an entire trial but could vary in color from trial-to-trial. It also served as the fixation point that subjects were to look at up until stimulus emergence.

When the stimulus appeared above the occluder, participants were required to move the real-time cursor to a central start location. The stimulus then dropped down the vertical channel, disappeared behind the occluder, and then reappeared 1000 ms later in a continuous oblique-downward motion from the right or left outlet.

The outlets were approximately 20 cm lateral to and slightly above the central start location. Participants were instructed to initiate a rapid goal-directed pro- or anti-reaching movement as soon as the stimulus reappeared below the occluder. They were allowed to reach through the stimulus, i.e. they did not have to stop at the location of the stimulus. At the end of each trial, the following feedback was presented on the screen the inter-trial interval:

“HIT” ➔ when they correctly intercepted the stimulus on pro-reach trials or correctly moved away from the stimulus on anti-reach trials

“WRONG WAY” ➔ when the movement went into the wrong direction relative to the given instruction

“MISS” ➔ when the movement went into the correct direction but the stimulus was not intercepted with sufficient accuracy

After having completed the trial, the participant returned to the starting position and the subsequent trial was initiated. Participants completed 600 trials in total, divided into 6 separate blocks with 100 trials each. Stimulus location (left/right) and reaching instruction (pro/anti) were randomized on each trial. Postural position (stable chair/wobble stool) was counterbalanced across blocks. Thus, there were 75 repeats of each unique trial type (e.g., right pro-reaches from the stable chair).

We manipulated postural demands as follows. In the low postural demand condition, participants were seated on an office chair that was locked in such a way that swiveling and rotation were not possible (Figure 1B). Participants were instructed to place their feet on the ground wide apart, which further increased postural stability. In the high postural demand condition, participants were seated on a balance stool (Figure 1B). This stool had a rounded base, which required continuous stabilization to maintain an upright posture. In addition, participants placed their feet at shoulder width and on top of two tennis balls. This was done to further decrease the level of support that the feet provide, requiring stronger activation of upper body muscles for stabilization. In both postural positions, participants were instructed to not lean the head forward against the headrest that is part of the KINARM platform, not to lean back into the backrest, and not to hold onto the table with the left hand. These instructions were given with the intention to keep the general seating position as similar as possible between the two postures.

### Data processing and analysis

All analyses were completed using MATLAB (version R2019a, MathWorks Inc., Natick, Massachusetts, United States of America). On each trial, a white stimulus that was placed under a photodiode and was unseen by the participant appeared simultaneously with stimulus emergence below the occluder. All kinematic and EMG data were aligned to diode onset.

#### Kinematic analyses

Inclusion and exclusion criteria for trials were determined using the kinematic output from the KINARM manipulandum. Reaction time was defined as the time between stimulus reappearance until reach initiation. Reach initiation was identified as the moment at which the tangential peak velocity of the hand surpassed 5% of the peak velocity of that trial. Peak reach velocity was derived from the reach-related maximum tangential velocity of the hand between movement onset and movement offset, which is defined as the point at which the tangential velocity reached zero following the movement. Trials were labelled as ‘wrong way’ and excluded from all kinematic and EMG analyses if at any point following target onset the participant’s hand moved opposite of the instructed reach direction (i.e. away from the target on pro-reach trials, towards the target on anti-reach trials). Trials were labelled as ‘too fast’ and excluded if the participant’s reaction time was shorter than 130 ms, indicating guesses or anticipatory movement. This is in line with earlier research (Gilchrist et al., 2024). We also excluded trials with reaction times that surpassed 500 ms, indicating distraction or inattention. In addition, kinematic data from all trials were manually checked via customized MATLAB GUIs that permitted the exclusion of clearly atypical trials (e.g. anticipatory movements prior to stimulus reappearance, or multi-step reaches).

We further investigated the variability of the KINARM’s handle position at baseline in the 200ms prior to target onset. The purpose of this analysis was to investigate if the unstable postural condition significantly increased instability, which may be reflected in an increased variability in handle position at baseline as participants may have used the handle to stabilize their posture. Because visual inspection of the variance revealed that the data was right-skewed, we used a median absolute deviation (MAD) metric to determine differences in variability between conditions, as, unlike the standard deviation, the MAD provides a more robust estimate of variability by relying on the median rather than the mean. This reduces the impact of outliers and ensures a more accurate representation of baseline variability. For each trial within a postural condition (stable/unstable), we calculated the variance of the position data during the 200 ms before target onset, using only ‘correct’ trials. We then determined the median variance across trials for either postural condition. Next, we computed the MAD by calculating the median of the absolute differences between each trial’s variance and the median variance within that condition. A cutoff threshold for potential outliers was defined as three times the MAD around the median variance and any variance values exceeding this threshold were removed. Finally, we averaged the remaining variances, yielding a single variance metric for each postural condition per subject.

#### EMG analyses

Wherever possible, we used EMG data recorded from the sternal head of the pectoralis muscle. Recordings from the clavicular head of pectoralis were used only if the sternal recording was of low quality (e.g., the signal-to-noise ratio was clearly better on the clavicular recording, or if the recording electrodes on the sternal head became loose during the experiment). We used the sternal recording in 95% of the younger participants (21/22) and in 92% of the elderly participants (12/13). EMG data were amplified by 1000, sampled at 1kHz, and bandpass filtered between 20 and 450 Hz by the Bagnoli-8 system. Offline, EMG signals were full-wave rectified and a 3-point smoothing function was applied. Comparisons of the magnitude of recruitment for the younger versus elderly subjects required that we first normalize EMG data to the level of recruitment in the 200 ms preceding the stimulus emergence, when participants were holding their arms stable against the background load.

#### EVR presence, latency, and magnitude

The detection of EVRs was performed using previously established methodologies (Corneil et al., 2004; Pruszynski et al., 2010) that use a time-series receiver operating characteristic (ROC) analysis. Independent time-series ROC analyses were conducted for each task variant (e.g., pro-reaches from the unstable condition). This approach assesses the comparative distribution of muscle recruitment at each time point from 100 ms before to 200 ms after stimulus onset for trials where the stimulus appeared on the left versus the right, deriving the area-under-the-curve (AUC) metric which represents the probability of correctly distinguishing stimulus location based solely on EMG activity. An AUC of 0.5 reflects chance-level discrimination, whereas values of 1.0 and 0.0 indicate fully accurate or fully inaccurate discrimination, respectively. An EVR was deemed to be present if the AUC value crossed and stayed above a threshold of 0.6 for at least 8 out of the following 10 samples within the time window of 80 to 120 ms post-stimulus. This point determined the *discrimination latency* of the EVR. Discrimination latency determined in this way may be confounded by the vigor of the EVR response, as stronger responses may climb to threshold more rapidly than weaker responses; this may be particularly problematic given past evidence that elderly subjects have slower and less vigorous on-line corrections (Kimura et al., 2015; O’Rielly & Ma-Wyatt, 2020; Sarlegna, 2006). Given this, we also employed an alternative approach that identifies the inflection point where the ROC curve first deviates from chance toward the discrimination threshold (AUC = 0.6). This was done by fitting a DogLeg regression to the ROC curve at each millisecond from 50 ms before stimulus onset up to the EVR latency (as previously employed by Contemori et al., 2021a; Goonetilleke et al., 2015). The EVR *onset latency* was determined as the later of two candidate time points: (1) the point minimizing the squared error between the ROC curve and the regression fit, or (2) the final local minimum in the ROC curve before the discrimination latency. The EVR latencies from both methods will be reported in the Results section. The EVR magnitude was calculated as the integral of the positive difference in mean EMG activity between leftward and rightward stimulus emergence over the 30 ms after the discrimination latency in the stable posture.

As part of our investigation of age-related effects on EVR expression, we were also interested in the ability to modulate EVR magnitudes depending on the instruction to prepare for a Pro-reach vs. Anti-Reach trial. We therefore derived a Modulation index (as previously reported by Gilchrist et al., 2024), which was calculated as the difference in EVR Magnitude on Pro-reach vs. Anti-reach trials, divided by the sum of the magnitudes on Pro-reach and Anti-reach trials. The Modulation index ranges from −1 to 1. A value of 1 indicates strong pro-reach EVRs and weak anti-reach EVRs, a value of 0 means equal magnitude EVRs on Pro-reach and Anti-reach trials, whereas a value of −1 indicates strong EVRs on anti-reach trials but not on pro-reach trials.

#### Statistical analysis

Statistical analyses were performed using MATLAB (version R2019a). The level of significance was set to *p <* .05 for all analyses. Repeated Measures ANOVAs were used to investigate the effects of group (Young vs. Elderly), posture (Stable vs. Unstable) and instruction (Pro-reach vs. Anti-reach) on error rates, reaction times, maximum reach velocities, and EVR magnitudes. To compare EVR prevalence between the HC and PD groups we used Fisher’s exact test. Because some participants did not exhibit EVRs in certain conditions (e.g., anti-reaches), we used linear mixed models to investigate EVR latencies. Unlike ANOVAs, LMMs do not rely on list-wise deletion for missing data, allowing us to maximize statistical power and minimize bias in our analysis. We defined group, posture and instruction as fixed effects and participant ID as a random effect.

## Results

### Influence of postural manipulation on handle position prior to target emergence

First, we examined whether our manipulation of postural demands had the intended effect on postural instability, as inferred from the variability of the position of the KINARM handle prior to target emergence. The overall variance of the position data did not differ significantly between young and elderly participants (*t(23.21)* = 1.86, *p =* .08), but consistent with the rationale of our postural challenge both groups had a significantly higher variability in the unstable condition (elderly: *M =* 3.3e-08 m, *SD =* 1.6e-08 m, *t(12)* = −7.47, *p <* .001; young: *M =* 7.0e-08 m, *SD =* 8.8e-08 m, *t(21)* = −3.7, *p =* .0012) compared to the stable condition (elderly: *M =* 3.3e-09 m, *SD =* 2.0e-09 m; young: *M =* 6.3e-09 m, *SD =* 8.8e-09 m).

### Fewer errors, longer reaction times, and slower reach velocities in the elderly

Figure 2 displays the results for the various behavioral measures of the reaching movement in young and older participants. Generally, error rates were low, with the overall proportion of reaches in the wrong direction being less than ∼10-15% (Figure 2A). Interestingly, the elderly participants (*M =* 7.7%, *SD =* 6.9%) made significantly fewer errors compared to younger participants (*M =* 13.8%, *SD =* 9.1%; *Group*, *F*(1,269) = 40.23, *p <* .001), but postural demand did not have an effect on wrong-way error rates (*Posture, F*(1,269) = 1.06, *p =* .30). Fewer errors were made on pro-reach trials (*M =* 8.1%, *SD =* 6.5) compared to anti-reach trials (*M =* 15%, *SD =* 9.6%; *Instruction*, *F*(1,269) = 48.71, *p <* .001). There were no significant interaction effects on error rates.

**Figure 2.**
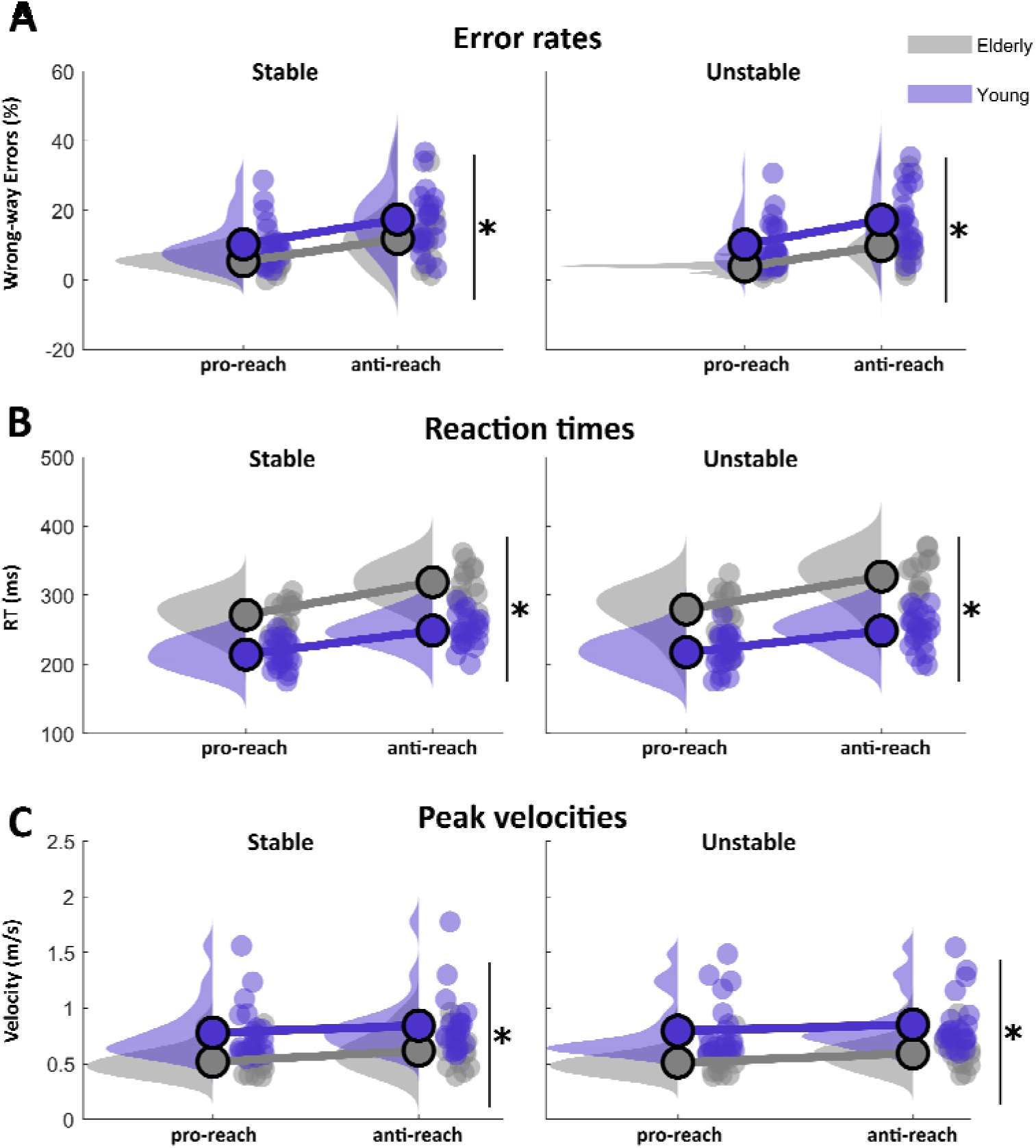
Behavioral results. The dots indicate individual data points of participants from the young (purple) and from the elderly (grey) group. The density plots indicate the distribution of the data. Asterisks show significant group differences (p < .05).

In terms of reaction time (Fig. 2B), the younger participants exhibited significantly shorter reaction times (*M =* 249ms, *SD =* 27ms) compared to the elderly participants (*M =* 280ms, *SD =* 25ms; *Group*, *F(*1) = 389.24, *p <* 0.001), independent of posture (*Group × Posture*, *F*(1,269) = 0.96, *p =* .33). As expected, reaction times were significantly shorter on pro-reach trials (*M =* 242ms, *SD =* 21ms) compared to anti-reach trials (*M =* 287ms, *SD =* 18ms; *Instruction, F*(1,269) = 139.91, *p <* 0.001).

Finally, we also analyzed the peak velocity of the reaching movement (Fig. 2C), and found that younger participants achieved higher reach velocities (*M =* .82 m/s, *SD =* .26 m/s) compared to the elderly group (*M =* .56 m/s, *SD =* .15 m/s, *Group* (*F(1,269)* = 85.49, *p <* .001). Somewhat surprisingly, reach velocities on Pro-reach trials were significantly slower (*M =* .69 m/s, *SD =* .25 m/s) compared to Anti-reach trials (*M =* .76 m/s, *SD =* .26 m/s; *Instruction, F*(1,269) = 7.33, *p =* .007). Recall however that the emerging stimulus (or its imagined location on anti-reach trials) moves in an oblique down-and-out direction upon emergence, hence the longer reaction times on anti-reach trials are likely associated with larger reaching movements, leading to higher peak velocities. Indeed, movement amplitudes (i.e. the distance between starting position and end position of the reach) were significantly larger on anti-reach trials (*M* = 16 cm, *SD* = 2 cm) compared to pro-reach trials (*M* = 14 cm, *SD* = 1 cm; *F*(1,273) = 81.8, *p* < .001). There was no effect of posture on peak velocities (*p =* .95).

Overall, this pattern of elderly participants having fewer errors, longer reaction times and slower peak velocities compared to younger participants points towards a potential speed-accuracy trade-off, with the elderly participants erring more on the side of caution (i.e. prioritizing accuracy), independent of postural condition.

### Muscle recruitment in two representative subjects during pro- and anti-reaches

We now discuss the patterns of muscle recruitment following stimulus presentation. Figure 3 shows muscle recordings of the right pectoral major from a younger (upper half) and elderly participant (lower half), respectively. The data from these two subjects demonstrate some of the key features of muscle recruitment.

**Figure 3.**
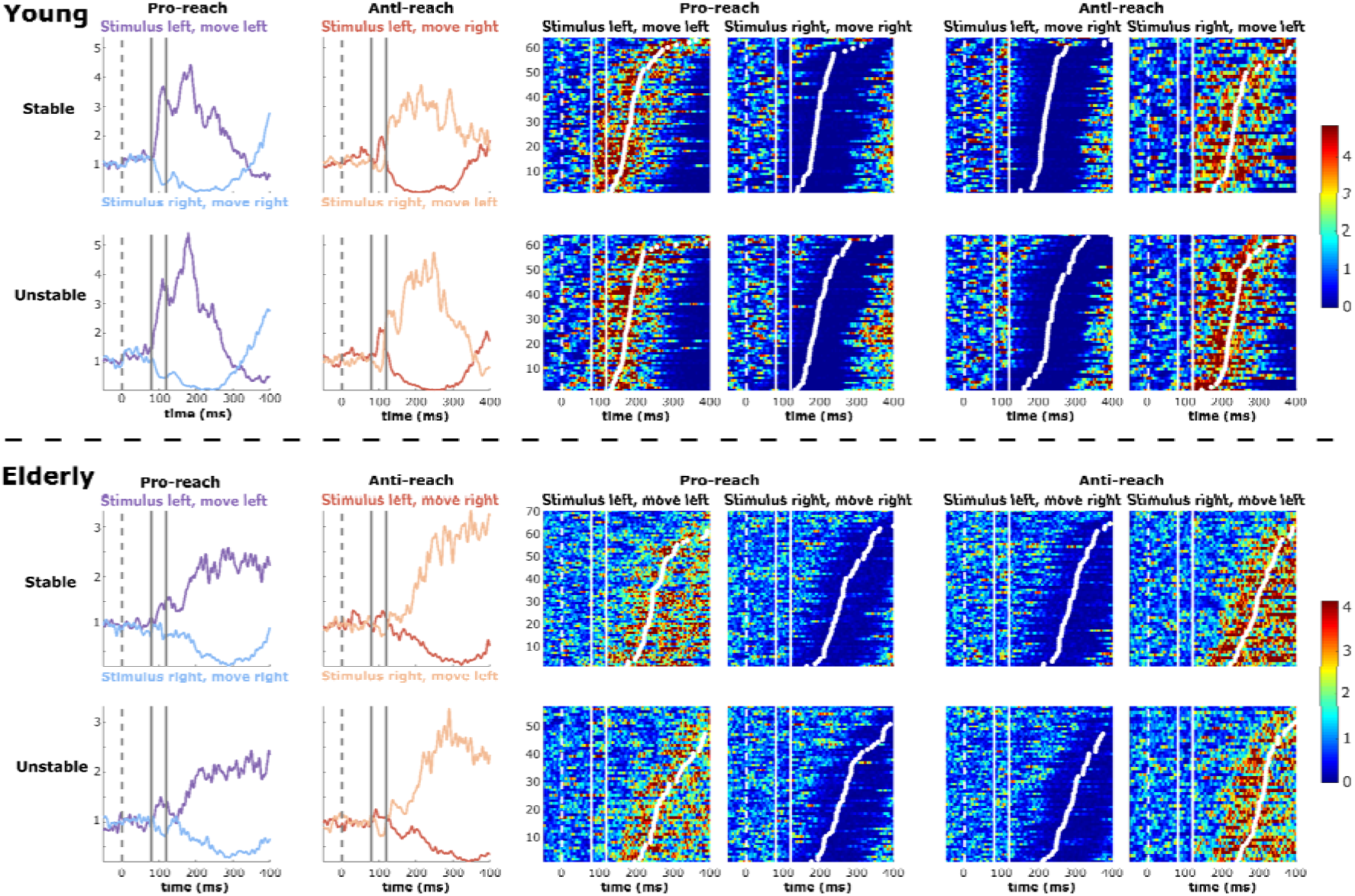
Muscle activity from two exemplar subjects. First two columns show averaged EMG data from right pectoralis across pro-reach trials (1^st^ column) and anti-reach trials (2^nd^ column) from a stable (upper row) and unstable position (lower row), normalized to baseline activity. The EVR activity is highlighted by the vertical solid lines. Note how EVRs increase following left target presentation, even if the instruction was to reach away from the target (on anti-reach trials). Columns 3-6 show heatmaps of the single trial EMG data. Activity is aligned to target onset (vertical dashed white line), EVR activity is highlighted by the solid white lines. Data is sorted by reaction times (white dots).

The first two columns depict the mean EMG activity across trials on pro-reach trials and on anti-reach trials for the stable (first row) and unstable posture (second row). The remaining columns show the trial-by-trial data of muscle recruitment, aligned to stimulus emergence. Focusing first on pro-reaches towards the left (purple), there was an initial burst of muscle activity at ∼75ms in both the stable and unstable postural condition, which is the express visuomotor response (the time window is indicated by the two vertical lines).

As is visible in the trial-by-trial data (colored heatmaps in third column), the EVR is more time-locked to stimulus presentation that reach movement onset, but larger EVRs tend to precede shorter reaction times, which is in line with previous findings (Pruszynski et al., 2010; Wood et al., 2015). The general pattern of muscle recruitment is very similar in our exemplar young and elderly participants, but both EVR magnitudes and the subsequent muscle recruitment were much more vigorous in the younger participant, which presumably relates to the shorter reaction times in the younger participant (white dots). In both participants, there are no clearly discernable differences between muscle recruitment on pro-reaches from a stable versus an unstable posture.

On anti-reach trials, EVR expression was tied to the side of stimulus emergence, with clear EVRs when targets were presented on the left, even if the reaching movements itself were correctly made towards the right (and vice versa). This is in line with prior research (Gu et al., 2016; Kozak et al., 2020). Again, the general pattern of muscle recruitment on anti-reach trials is similar across subjects, but EVR responses seemed more vigorous in the younger subject. In both the young and elderly exemplar subject, the magnitude of the EVR is clearly muted on anti-compared to pro-reach trials.

### EVRs less prevalent in elderly participants

EVRs were robustly detected during pro reaches in both groups. During pro-reach trials towards the left, all younger participants exhibited EVRs (22/22; 100%) in both postural conditions, whereas only 10 out of 13 (77%) elderly participants had EVRs in either postural condition (9 of them with EVRs in both conditions). This difference in EVR prevalence was significant between the two groups (*p =* .02). During anti-reach trials in the stable condition, we identified EVRs in 8 younger participants (36%) and 4 elderly participants (31%). In the unstable condition, these prevalences were 7 (32%) and 3 (23%) for the two groups respectively. The difference between groups was not significant (*p =* .58).

### Weaker EVRs in elderly participants

In line with the observations from the two representative subjects described above, normalized EVR magnitudes were smaller in the elderly participants (*M =* 8.6 a.u., *SD =* 6.5 a.u.) compared to the younger participants (*M =* 24.1 a.u., *SD =* 23.8 a.u.; *group, F(1,269)* = 67.7, *p <* .001); the difference between groups was larger within pro-reach trials (elderly: *M =* 33.6 a.u., *SD =* 21.1 a.u.; young: *M =* 43.4 a.u., *SD =* 18.7 a.u.), compared to anti-reach trials (elderly: *M =* 3.8 a.u., *SD =* 3.4 a.u.; young: *M =* 4.8 a.u., *SD =* 5.3 a.u.; *group × instruction* (*F*(1,269) = 24.76, *p <* .001, see Figure 4A). Overall, EVR magnitudes were significantly smaller on anti-reach trials (*M =* 4.4 a.u., *SD =* 4.7 a.u.) compared to pro-reach trials (*M =* 33.4 a.u., *SD =* 21.1 a.u., *instruction* (*F*(1,269) = 54.26, *p <* .001). There was no effect of posture on EVR magnitudes (*p =* .94).

**Figure 4.**
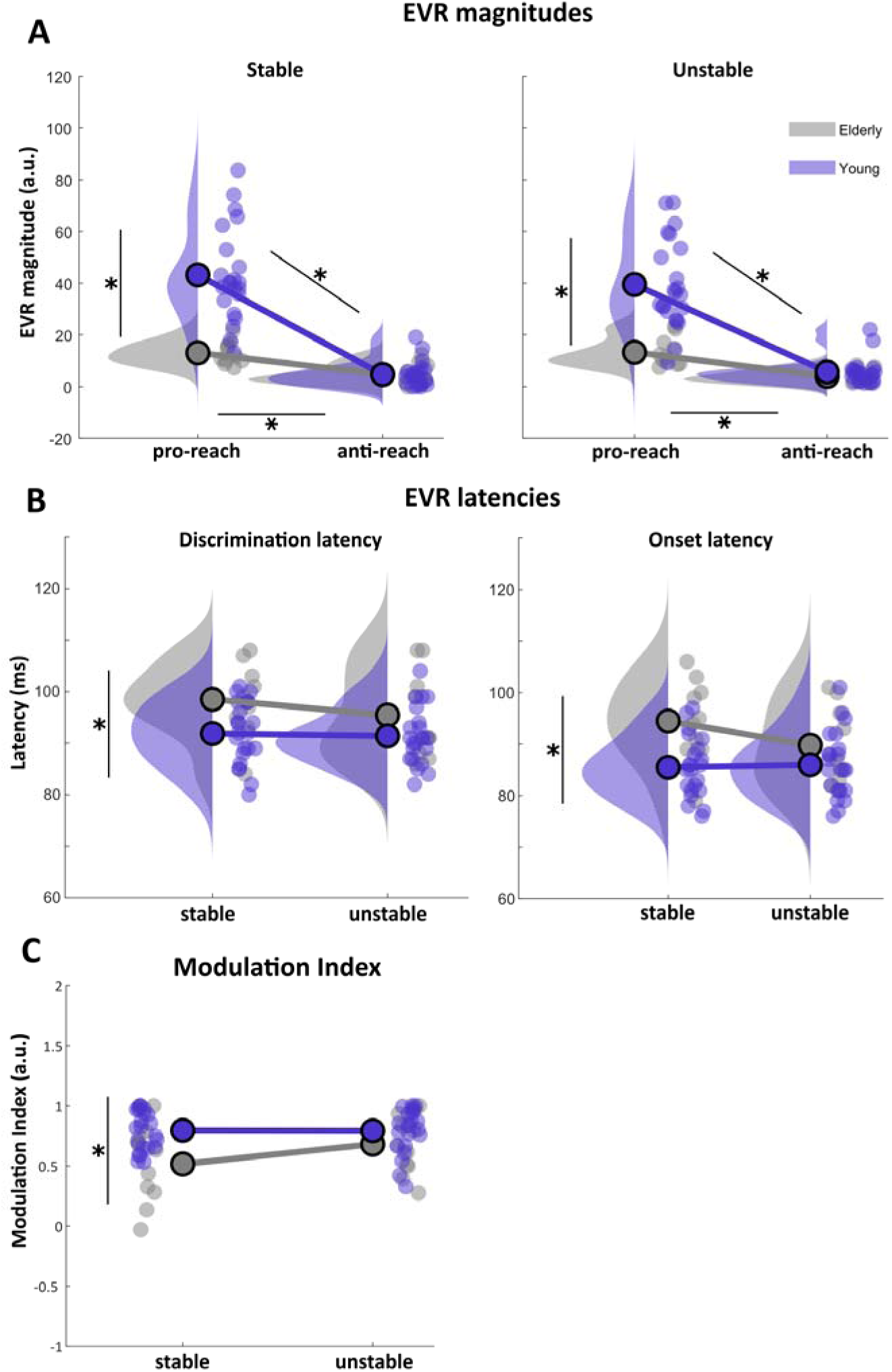
EVR-related outcomes. The dots indicate individual data points of participants from the young (purple) and from the elderly (grey) group. The density plots indicate the distribution of the data. The EVR latencies show the discrimination latency (crossing of the AUC threshold of 0.6, B, first panel) and the onset latency (DogLeg regression method, B, second panel). Note the absence of a density plot for the Modulation index (C). The modulation index is capped between −1 and + 1 and a density plot would not adequately reflect these bounds. Asterisks show significant effects (p < .05) of group (vertical lines), instruction (horizontal lines) and group × instruction interaction (diagonal lines).

### Delayed EVRs in elderly participants

As described in the Methods, we used two measures to characterize the timing of the EVR. First, we defined the first instance that the time-series ROC curve surpassed a threshold of 0.6 (Fig. 4B, first panel). With this measure, discrimination latencies were significantly earlier in the younger participants (*M =* 91 ms, *SD =* 5.1) compared to the elderly participants (*M =* 98 ms, *SD =* 8 ms; *Group,* β = 6.8, *SE* = 2.3, *p =* .003, *95% CI* [2.27 11.39]), independent of posture (*Group × Posture*, β = −2.9, *SE* = 3.25, *p =* .37, *95% CI* [-9.4 3.5], see Figure 4B). Discrimination latencies were significantly earlier on pro-reach trials (*M =* 93 ms, *SD =* 6.9 ms) compared to anti-reach trials (*M =* 97 ms, *SD =* 6.8 ms; *Instruction*, β = 4.6, *SE* = 2.21, *p =* .04, *95% CI* [0.2 9]).

Second, and guided by a potential confound of this discrimination latency with response vigor, as reflected in the increased EVR magnitudes in the younger participants, we also used a DogLeg regression method. This method detects the timepoint at which the time-series ROC begins to increase, and as expected, this timepoint occurs earlier (by ∼6 ms) than the time at which the time-series ROC exceeds the threshold value of 0.6. The effects of age and postural condition were, however, similar, as was the magnitude of the time difference in EVRs in the younger vs elderly participants: younger participants had earlier onset latencies (*M =* 85ms, *SD =* 5.1ms) compared to the elderly participants (*M =* 93ms, *SD =* 7.6ms; *Group,* β = 9.0, *SE* = 2.53, *p <* .001, *95% CI* [3.98 14.0]), independent of posture (*Group × Posture*, β = −5.3, *SE* = 3.63, *p =* .15, *95% CI* [-12.5 1.9], Figure 4B, second panel). Using this method, there was no effect of instruction on onset latencies (β = 3.5, *SE* = 2.4, *p =* .15, *95% CI* [-1.3 8.4]). Thus, older participants exhibited longer latency EVRs, regardless of whether one characterizes the timepoint at which the time-series ROC curve begins to increase, or the timepoint at which the curve exceeds a threshold.

### Higher Modulation index in younger participants

The modulation index was calculated to investigate the participant’s ability to contextually modulate EVR magnitude depending on the instruction to reach towards or away from the target. Younger participants had a significantly higher modulation index (*M* = 0.79, *SD* = 0.16) compared to the elderly subjects (*M* = 0.60, *SD* = 0.29; *F*(1,269) = 12.16, *p* < .001), but this effect seems to be driven by the weaker pro-reach EVRs in the group of elderly participants (Figure 4C). There were no main or interaction effects of posture (p ≥ 0.14).

## Discussion

We investigated express visuomotor responses (EVRs) in the upper limbs of healthy young and elderly participants during goal-directed reaches from a stable and an unstable sitting posture. Our two objectives were to investigate age-related effects on EVR expression and to investigate the interaction between postural control and EVR expression in the upper body. In younger participants, EVRs were robustly present, with no significant differences in their prevalence, timing, or magnitude across postural conditions. In comparison, elderly participants exhibited less frequent, delayed, and weaker EVRs, as well as longer movement reaction times and slower peak velocities compared to younger participants. As with younger participants, these measures did not differ between postural conditions.

### Age-related effects on EVR expression

This study is the first to directly compare the expression of EVRs between younger and elderly participants in the exact same behavioral paradigm. Our findings revealed clear differences between the two groups: although most elderly participants exhibited EVRs, their prevalence was overall lower, latencies were longer and magnitudes were smaller. Importantly, our use of the dog-leg regression affirms that the increased EVR latencies in the elderly are not just the result of decreasing EVR magnitude; EVRs started ∼7 ms later in the elderly regardless of the analytical approach. Behaviorally, elderly participants also had longer reaction times and slower peak velocities compared to younger participants. The notable differences in performance between younger and elderly participants raise several important questions: Why are EVRs less prevalent, weaker, and slower in the elderly? Are these differences purely a result of age-related declines in neural pathways, or do they also reflect changes in motivation or strategy?

The effect of aging we observed on the timing of the fast visuomotor network is reminiscent of age-related slowing of express saccades (Munoz et al., 1998; O’Rielly & Ma-Wyatt, 2020; Peltsch et al., 2011). EVRs are proposed to be the skeleto-motor homologue of express saccades (Corneil Munoz reference), which rely on the integrity of the superior colliculus (Schiller et al., 1987). Similarly during reaching, elderly participants initiate mid-reach corrections in response to sudden target jumps later or not at all compared to younger subjects (Kimura et al., 2015; O’Rielly & Ma-Wyatt, 2020). In the lower body, studies on obstacle avoidance, which involve rapid step adjustments to the ongoing stepping movement in response to sudden obstacles, demonstrated longer reaction times and longer muscle response onsets in elderly participants (Weerdesteyn et al., 2005, 2007). Such delayed responses during online corrections, both kinematically and on a muscular level, are consistent with a delayed and muted EVR (Mekhaiel et al., 2025). Collectively, these findings highlight age-related declines in rapid motor responses across multiple domains.

What may be the cause of these age-related declines across various behaviors that, just like EVRs, may all rely on the same underlying network? One possible explanation is age-related changes in neural and motor systems at various levels of the neuraxis. These changes can begin as early as the eye, where common aging-related changes like cataract, reduced pupil dilation, or presbyopia may affect how visual information is registered and subsequently processed for input into the fast visuomotor network. Since sensory properties of visual targets, such as stimulus contrast or spatial frequency influence EVR expression (Kozak et al., 2019; Kozak & Corneil, 2021), it is not surprising that deficits in stimulus processing may lead to less frequent, weaker, and delayed EVRs in elderly participants. Similarly, the subcortical tecto-reticulospinal pathway that is thought to project EVRs from superior colliculus to the motor periphery may also become less efficient with age, leading to delayed and weakened outputs. Supporting this, recent studies have demonstrated age-related decreases in the excitability of the reticulospinal tract (Mooney & Celnik, 2025), as well as slower overall conduction velocities to the motor periphery (Palve & Palve, 2018).

Another factor to consider is the potential difficulty in motivating elderly participants to perform tasks requiring rapid, reflexive movements. Unlike younger participants, and in line with previous research, elderly individuals might prioritize caution and accuracy over speed, particularly in tasks that could be perceived as posturally challenging (Salthouse, 1979; Starns & Ratcliff, 2010).

While such caution can be adaptive when balance is genuinely challenged, the delayed and slower movements observed in the stable postural condition suggest that older adults may be overly cautious even when it is unnecessary. During the pilot phase of a previous study on EVRs in stepping (Billen et al., 2024), we also observed that elderly participants found it harder to step fast as compared to younger participants (Billen et al., 2023). This reluctance to step fast was partly overcome by further emphasizing the importance of speed in our instructions, and repeating this instruction frequently during the experiment, yet in the elderly participants EVRs still appeared to be suppressed and RTs delayed (Billen et al., 2024). A similar mechanism may be at play in the current study, where elderly individuals might subconsciously prioritize stability and avoid situations that require rapid movements. Such cognitive strategies have generally been shown to affect EVR response vigor (i.e., the magnitude of the response) rather than the timing of the EVR (Contemori et al., 2021, 2022; Gu et al., 2016; Wood et al., 2015).

Together, impairments in visual processing, reduced efficiency of the reticulospinal pathway, and a possible subconscious tendency to prioritize safety over speed can all contribute to decreased adaptability in our dynamic environments, which may have significant implications for daily life. Such age-related changes may for example affect the ability to safely drive a car, which heavily relies on the ability to rapidly transform visual information into appropriate motor commands both in the upper body (e.g. for steering) and lower body (e.g. for braking). Similarly, declines in the ability to rapidly and flexibly adjust stepping movements may increase the risk of falls. Implementing interventions to maintain some degree of flexibility may be crucial in the aging population.

### Postural instability did not affect EVR expression in the upper limb in either group

Our findings showed no significant effects of posture on any of the main outcome variables. This result may partially stem from the characteristics of the wobble stool used to introduce postural instability. Anecdotally, some participants experienced mild instability during the first few trials as they acclimated to the wobble stool. However, this instability appeared to diminish relatively quickly. While our kinematic results indicate increased instability when using the wobble stool, this increase in postural challenge may not have been sufficient to affect expression of the fast visuomotor network.

Moreover, the role of core muscles likely contributed to the lack of observable effects. In the unstable condition, the core muscles were probably more engaged to stabilize the torso, which might have helped participants maintain a steady posture without restricting the activation of the pectoralis major involved in the reaching task. This raises an interesting question for future research, though recording high-quality surface EMG from core muscles presents challenges. Sitting compresses the trunk, complicating accurate electrode placement and potentially reducing signal quality. These issues may be further amplified in older adults, who often have increased adipose tissue, which can further lower the signal-to-noise ratio.

Additionally, the use of a handle as part of the KINARM setup may have inadvertently provided participants with a source of stability, further reducing the postural challenge. This is highlighted by the finding that the variability of the handle position at baseline increased in the unstable condition, indicating the while the wobble did induce more instability, the handle was also used in order to stabilize. Future paradigms could explore reaching tasks without added stability from a handle, creating a more demanding postural environment while still involving rapid goal-directed movements. This adjustment may better reveal the potential relationship between postural control and EVR expression in the upper body.

## Conclusion

This study is the first to compare EVR expression between younger and elderly participants, providing novel insights into age-related changes in rapid visuomotor responses. While most elderly participants still exhibited robust EVRs, they were weaker, less frequent, and delayed, indicating potential degradation of the fast visuomotor network with increasing age. These findings serve as an important reference for future research, particularly when considering elderly participants as a control group in studies investigating clinical populations, such as individuals with Parkinson’s disease.

Although we did not establish a clear interaction between postural control and EVR expression in the upper limbs, this does not preclude its existence. Previous research has demonstrated this interaction in stepping, and with adjustments to experimental design - such as increasing postural demands and incorporating more precise measurements of upper body stability - we may better capture its effects in upper limb movements. Future studies should refine methodologies to further explore how postural control influences rapid visuomotor responses in order to increase our understanding of rapid visuomotor transformations in both healthy aging and clinical populations.

## Author Contributions

*Lucas Billen:* Conceptualization, Methodology, Software, Formal analysis, Investigation, Data curation, Writing – Original Draft, Review & Editing, Visualization; *Madeline Gilchrist:* Methodology, Software, Formal analysis, Investigation, Writing – Review & Editing; *Brian Corneil:* Conceptualization, Methodology, Software, Resources, Writing - Review & Editing, Supervision, Funding acquisition; *Vivian Weerdesteyn:* Conceptualization, Methodology, Resources, Writing - Review & Editing, Supervision, Project administration, Funding acquisition

## Data availability statement

The authors confirm that the data supporting the findings of this study are available within the article and its supplementary materials.

## References

Billen, L. S., Corneil, B. D., & Weerdesteyn, V. (2023). Evidence for an Intricate Relationship Between Express Visuomotor Responses, Postural Control and Rapid Step Initiation in the Lower Limbs. Neuroscience. 10.1016/j.neuroscience.2023.07.025

Billen, L. S., Nonnekes, J., Corneil, B. D., & Weerdesteyn, V. (2024). Lower-limb express visuomotor responses are spared in Parkinson’s Disease during step initiation from a stable position (p. 2024.11.29.625631). bioRxiv. 10.1101/2024.11.29.625631

Bugnariu, N., & Sveistrup, H. (2006). Age-related changes in postural responses to externally- and self-triggered continuous perturbations. Archives of Gerontology and Geriatrics, 42(1), 73–89. 10.1016/j.archger.2005.05.003

Contemori, S., Loeb, G. E., Corneil, B. D., Wallis, G., & Carroll, T. J. (2021). The influence of temporal predictability on express visuomotor responses. Journal of Neurophysiology, 125(3), 731–747. 10.1152/jn.00521.2020

Contemori, S., Loeb, G. E., Corneil, B. D., Wallis, G., & Carroll, T. J. (2022). Symbolic cues enhance express visuomotor responses in human arm muscles at the motor planning rather than the visuospatial processing stage. Journal of Neurophysiology. 10.1152/jn.00136.2022

Corneil, B. D., & Munoz, D. P. (2014). Overt Responses during Covert Orienting. Neuron, 82(6), 1230– 1243. 10.1016/j.neuron.2014.05.040

Corneil, B. D., Olivier, E., & Munoz, D. P. (2004). Visual responses on neck muscles reveal selective gating that prevents express saccades. Neuron, 42(5), 831–841. 10.1016/s0896-6273(04)00267-3

Day, B. L., & Brown, P. (2001). Evidence for subcortical involvement in the visual control of human reaching. Brain, 124(9), 1832–1840. 10.1093/brain/124.9.1832

Day, B. L., & Lyon, I. N. (2000). Voluntary modification of automatic arm movements evoked by motion of a visual target. Experimental Brain Research, 130(2), 159–168. 10.1007/s002219900218

Fautrelle, L., Ballay, Y., & Bonnetblanc, F. (2010). Muscular synergies during motor corrections: Investigation of the latencies of muscle activities. Behavioural Brain Research, 214(2), 428–436. 10.1016/j.bbr.2010.06.015

Fautrelle, L., Prablanc, C., Berret, B., Ballay, Y., & Bonnetblanc, F. (2010). Pointing to double-step visual stimuli from a standing position: Very short latency (express) corrections are observed in upper and lower limbs and may not require cortical involvement. Neuroscience, 169(2), 697– 705. 10.1016/j.neuroscience.2010.05.014

Giesbers, I., Billen, L., van der Cruijsen, J., Corneil, B. D., & Weerdesteyn, V. (2025). Cortical dynamics underlying initiation of rapid steps with contrasting postural demands. Neuroscience, 575, 104–121. 10.1016/j.neuroscience.2025.04.026

Glover, I. S., & Baker, S. N. (2019). Multimodal stimuli modulate rapid visual responses during reaching. Journal of Neurophysiology, 122(5), 1894–1908. 10.1152/jn.00158.2019

Goonetilleke, S. C., Katz, L., Wood, D. K., Gu, C., Huk, A. C., & Corneil, B. D. (2015). Cross-species comparison of anticipatory and stimulus-driven neck muscle activity well before saccadic gaze shifts in humans and nonhuman primates. Journal of Neurophysiology, 114(2), 902–913. 10.1152/jn.00230.2015

Gu, C., Wood, D. K., Gribble, P. L., & Corneil, B. D. (2016). A Trial-by-Trial Window into Sensorimotor Transformations in the Human Motor Periphery. The Journal of Neuroscience, 36(31), 8273– 8282. 10.1523/JNEUROSCI.0899-16.2016

Kimura, D., Kadota, K., & Kinoshita, H. (2015). The impact of aging on the spatial accuracy of quick corrective arm movements in response to sudden target displacement during reaching. Frontiers in Aging Neuroscience, 7. 10.3389/fnagi.2015.00182

Kozak, R. A., Cecala, A. L., & Corneil, B. D. (2020). An Emerging Target Paradigm to Evoke Fast Visuomotor Responses on Human Upper Limb Muscles. JoVE (Journal of Visualized Experiments), 162, e61428. 10.3791/61428

Kozak, R. A., & Corneil, B. D. (2021). High-contrast, moving targets in an emerging target paradigm promote fast visuomotor responses during visually guided reaching. Journal of Neurophysiology, 126(1), 68–81. 10.1152/jn.00057.2021

Kozak, R. A., Kreyenmeier, P., Gu, C., Johnston, K., & Corneil, B. D. (2019). Stimulus-Locked Responses on Human Upper Limb Muscles and Corrective Reaches Are Preferentially Evoked by Low Spatial Frequencies. eNeuro, 6(5). 10.1523/ENEURO.0301-19.2019

Maki, B. E., & McIlroy, W. E. (1996). Postural Control in the Older Adult. Clinics in Geriatric Medicine, 12(4), 635–658. 10.1016/S0749-0690(18)30193-9

Mekhaiel, D. Y., Goodale, M. A., & Corneil, B. D. (2025). Are online corrections to visual targets really a distinct class of movement? (p. 2025.03.14.643316). bioRxiv. 10.1101/2025.03.14.643316

Mooney, R. A., & Celnik, P. A. (2025). Effector-dependent decline in strength and subcortical motor excitability with aging. Neurobiology of Aging, 147, 98–104. 10.1016/j.neurobiolaging.2024.12.008

Munoz, D. P., Broughton, J. R., Goldring, J. E., & Armstrong, I. T. (1998). Age-related performance of human subjects on saccadic eye movement tasks. Experimental Brain Research, 121(4), 391–400. 10.1007/s002210050473

Nonnekes, J. H., Talelli, P., de Niet, M., Reynolds, R. F., Weerdesteyn, V., & Day, B. L. (2010). Deficits Underlying Impaired Visually Triggered Step Adjustments in Mildly Affected Stroke Patients. Neurorehabilitation and Neural Repair, 24(4), 393–400. 10.1177/1545968309348317

O’Rielly, J. L., & Ma-Wyatt, A. (2020). Saccade dynamics during an online updating task change with healthy aging. Journal of Vision, 20(13), 2. 10.1167/jov.20.13.2

Palve, S. S., & Palve, S. B. (2018). Impact of Aging on Nerve Conduction Velocities and Late Responses in Healthy Individuals. Journal of Neurosciences in Rural Practice, 9(1), 112–116. 10.4103/jnrp.jnrp_323_17

Peltsch, A., Hemraj, A., Garcia, A., & Munoz, D. P. (2011). Age-related trends in saccade characteristics among the elderly. Neurobiology of Aging, 32(4), 669–679. 10.1016/j.neurobiolaging.2009.04.001

Pruszynski, J. A., King, G. L., Boisse, L., Scott, S. H., Flanagan, J. R., & Munoz, D. P. (2010). Stimulus-locked responses on human arm muscles reveal a rapid neural pathway linking visual input to arm motor output. European Journal of Neuroscience, 32(6), 1049–1057. 10.1111/j.1460-9568.2010.07380.x

Reynolds, R. F., & Day, B. L. (2005). Rapid visuo-motor processes drive the leg regardless of balance constraints. Current Biology, 15(2), R48–R49. 10.1016/j.cub.2004.12.051

Salthouse, T. (1979). Adult age and the speed-accuracy trade-off. Ergonomics, 22(7), 811–821. 10.1080/00140137908924659

Sarlegna, F. R. (2006). Impairment of online control of reaching movements with aging: A double-step study. Neuroscience Letters, 403(3), 309–314. 10.1016/j.neulet.2006.05.003

Schiller, P. H., Sandell, J. H., & Maunsell, J. H. (1987). The effect of frontal eye field and superior colliculus lesions on saccadic latencies in the rhesus monkey. Journal of Neurophysiology, 57(4), 1033–1049. 10.1152/jn.1987.57.4.1033

Starns, J. J., & Ratcliff, R. (2010). The effects of aging on the speed-accuracy compromise: Boundary optimality in the diffusion model. Psychology and Aging, 25(2), 377–390. 10.1037/a0018022

Weerdesteyn, V., Nienhuis, B., & Duysens, J. (2005). Advancing age progressively affects obstacle avoidance skills in the elderly. Human Movement Science, 24, 865–880. 10.1016/j.humov.2005.10.013

Weerdesteyn, V., Nienhuis, B., Geurts, A. C. H., & Duysens, J. (2007). Age-related deficits in early response characteristics of obstacle avoidance under time pressure. The Journals of Gerontology. Series A, Biological Sciences and Medical Sciences, 62(9), 1042–1047. 10.1093/gerona/62.9.1042

Wood, D. K., Gu, C., Corneil, B. D., Gribble, P. L., & Goodale, M. A. (2015). Transient visual responses reset the phase of low-frequency oscillations in the skeletomotor periphery. European Journal of Neuroscience, 42(3), 1919–1932. 10.1111/ejn.12976

Zhang, Y., Smeets, J. B. J., Brenner, E., Verschueren, S., & Duysens, J. (2021). Effects of ageing on responses to stepping-target displacements during walking. European Journal of Applied Physiology, 121(1), 127–140. 10.1007/s00421-020-04504-4

